# RBN-2397, a PARP7 Inhibitor, Synergizes with Paclitaxel to Inhibit Proliferation and Migration of Ovarian Cancer Cells

**DOI:** 10.1101/2024.08.20.608802

**Authors:** Alexandra N. Spirtos, Marwa W. Aljardali, Sridevi Challa, Sneh Koul, Jayanthi S. Lea, W. Lee Kraus, Cristel V. Camacho

## Abstract

**Objectives:** Mono(ADP-ribosyl)ation (MARylation), a post translational modification of proteins, is emerging as an important regulator of the biology of cancer cells. PARP7 (TiPARP), a mono (ADP-ribosyl) transferase (MART), MARylates its substrate α-tubulin in ovarian cancer cells, promoting destabilization of microtubules, cell growth, and migration. Recent development of RBN-2397, a potent inhibitor that selectively acts on PARP7, has provided a new tool for exploring the role of PARP7 catalytic activity in biological processes. In this study, we investigated the role of PARP7 catalytic activity in the regulation of ovarian cancer cell biology via MARylation of α-tubulin.

**Methods:** Ovarian cancer cell lines (OVCAR4, OVCAR3) were treated with RBN-2397 and paclitaxel, both separately and in combination. Western blotting and immunoprecipitation confirmed the effects of RBN-2397 on α-tubulin MARylation and stabilization. Cell proliferation and migration were assessed, and α-tubulin stabilization was quantified using immunofluorescent imaging. RNA-sequencing was performed to assess the effects on gene expression changes.

**Results:** RBN-2397 inhibited PARP7 activity, decreasing α-tubulin MARylation, leading to its stabilization, and reducing cancer cell proliferation and migration. The addition of paclitaxel further enhanced these effects, highlighting a synergistic interaction between the two drugs. Mutating the site of PARP7-mediated MARylation on α-tubulin similarly resulted in microtubule stabilization and decreased cell migration in the presence of paclitaxel.

**Conclusions:** This study demonstrates that targeting PARP7 with RBN-2397, particularly in combination with paclitaxel, offers an effective strategy for inhibiting aggressive ovarian cancer cell phenotypes. Our findings underscore the potential of combining PARP7 inhibitors with established chemotherapeutics to enhance treatment efficacy in ovarian cancer.

## Introduction

Ovarian cancer is the second leading cause of death from gynecologic cancers, with around 20,000 new cases and 13,000 deaths estimated in 2024 [1]. While the 5-year survival has increased over the past few decades, it remains low at 50% [2]. Up front treatment focuses on cytoreductive surgery with platinum- and taxane-based chemotherapy; unfortunately, most patients will recur. Recurrent ovarian cancer has a poor prognosis, particularly in the platinum resistant cohort. Paclitaxel is often selected as the first-line treatment for platinum-resistant recurrent disease. However, the duration of response is typically less than 6 months [3]. Paclitaxel binds and stabilizes β-tubulin blocking microtubule depolymerization, which prevents mitosis and induces cell death [4]. Unfortunately, various mechanisms of paclitaxel resistance in ovarian cancer have been proposed [5]. Because of the limited treatment options for platinum resistant disease, any potential new targets are valuable. Improving response rates by leveraging the known mechanism of paclitaxel and by increasing paclitaxel sensitivity is one way to improve survival in the setting of recurrent ovarian cancer.

ADP-ribosylation is a post-translational modification that involves transfer of ADP-ribose from β-nicotinamide adenine dinucleotide onto target proteins, DNA, or RNA [6]. Proteins can be ADP-ribosylated with a single or mono ADP-ribose (MARylation) or with multiple ADP-ribose moieties (PARylation) [7]. Poly(ADP-ribose) polymerases (PARPs) are a family of enzymes that catalyze these modifications [6]. PARP inhibitors have been approved for use in the setting of ovarian cancer [8]. However, these clinical PARP inhibitors target primarily PARP1 and PARP2, which are involved in nuclear PARylation, yet the majority of PARPs (PARP 3, 4, 6-12, and 14-16) catalyze MARylation. Despite this, to date few mono (ADP-ribosyl) transferase (MART) enzymes have been harnessed for clinical uses [8].

PARP7 is a nuclear and cytosolic MART that functions in viral responses, gene regulation, and cytoskeleton regulation [7]. Studies have suggested that the targets of PARP7 include PARP7 itself [9], AHR [9], type I interferon response regulators [9], histones [10], and Liver X Receptors [11]. PARP7 also plays a role in cancer biology. Using The Cancer Genome Atlas (TCGA) database, we previously established that ovarian cancer tissues have relatively low levels of PARP7 mRNA compared to normal ovarian tissue. However, the malignant ovarian cancer cells specifically have higher PARP7 mRNA levels than any of the normal ovarian cell types [12]. OVCAR4 cells with siRNA-mediated PARP7 depletion were shown to have decreased growth, migration, and invasion in vitro, compared to control cells. RNA sequencing data revealed that the PARP7 depleted cells are enriched for genes involved in cell-cell adhesion, cell cycle arrest, apoptosis, and gene regulation, supporting the role of PARP7 in cell growth and motility [12]. Finally, we reported that PARP7 MARylates and destabilizes α-tubulin. Mutating the specific site of MARylation on α-tubulin blocks the microtubule destabilization, supporting α-tubulin MARylation by PARP7 as a new actionable therapeutic pathway for PARP7 inhibitors [12]. Recently, the selective PARP7 small molecule inhibitor RBN-2397 has been shown to have antitumor effects in lung cancer xenograft studies in part through activation of a type I interferon response [9]. Herein, we explore the biological effects of RBN-2397 in ovarian cancer and provide evidence for a novel combination effect between this PARP7 inhibitor and an existing chemotherapeutic agent, paclitaxel. Our findings suggest a promising new strategy for the treatment of aggressive ovarian cancers.

## Materials and Methods

### Cell culture

OVCAR3 and OVCAR4 serous ovarian cancer cell lines were purchased from the American Type Culture Collection (OVCAR3: ATCC; RRID: CVCL_0465, OVCAR4: ATCC; RRID: CVCL_1627). They were maintained in RPMI media (Sigma-Aldrich, R8758) supplemented with Glutamax, 10% fetal bovine serum and 1% penicillin/streptomycin. HEK 293T, normal embryonic kidney human cell line was purchased from ATCC (RRID: CVC_0063). All cell lines were authenticated for cell type identity using the GenePrint 24 system (Promega, B1870), and confirmed as *Mycoplasma*-free every 6 months using the Universal Mycoplasma Detection Kit (ATCC, 30-1012K). Fresh cell stocks were regularly replenished from the original stocks every few months (no more than 15 passages).

### Antibodies and drugs

The antibodies used were as follows: FLAG (Sigma-Aldrich, F3165; RRID: AB_259529), α-tubulin (Santa Cruz, sc-8035; RRID: AB_628408), TiPARP (PAPR7) (Thermo Fisher Life Technologies, PA5-40774; RRID: AB_2607074), β-tubulin (Abcam, ab6046; RRID: AB_2210370), and HA (Sigma-Aldrich, H3663; RRID: AB_262051). The custom recombinant antibody-like MAR binding reagent (anti-MAR) was generated and purified in-house (now available from Millipore Sigma, MABE1076; RRID: AB_2665469) [13]. Secondary antibodies for immunofluorescence included Alexa Fluor 594 goat anti-mouse IgG (ThermoFisher, A-11005; AB_2534073). Drugs included paclitaxel (MedChemExpress, HY-B0015) and RBN-2397, which was synthesized by Acme Bioscience, Inc.

### Dose response and cell proliferation assays

For the RBN-2397 and paclitaxel dose response assays, OVCAR4 and OVCAR3 were seeded at density of 1×10^4^ cells per well in 12 well plates. Increasing doses of RBN-2397 and paclitaxel were applied, respectively, and media with fresh drug was replenished every 2 days. Plates were collected and fixed at day 6. Cell proliferation was assessed using a crystal violet staining assay. The cells were grown to ∼50% confluence (approximately 1 day of growth) and then treated with RBN-2397, paclitaxel, or combination. The cells were collected every 2 days after treatment. After collections, the cells were washed with PBS, fixed for 10 min with 4% paraformaldehyde at room temperature, and stored in 4°C until all time points had been collected. The fixed cells were stained with 0.1% crystal violet in 20% methanol solution containing 200 mM phosphoric acid. After washing to remove unincorporated stain, the crystal violet was extracted using 10% glacial acetic acid and the absorbance was read at 595 nm. All growth assays were performed a minimum of three times using independent plating of cells to ensure reproducibility.

### Cell migration assays

Boyden chamber assays were used to determine the effects of various treatments on cell migration. One hundred thousand cells (for OVCAR4) and 5×10^4^ cells (for OVCAR3) were seeded into the migration chambers (Corning 353097) following the manufacturer’s protocols. Cells were trypsinized, resuspended in serum-free media, and collected by centrifugation at 300 g for 3 min at room temperature. The cell pellets were then resuspended in serum-free media and 500 μL of media containing 1×10^5^ OVCAR4 cells or 5×10^4^ OVCAR3 cells were plated into the top chamber along with the indicated treatments. The chambers were then incubated in serum-containing media with FBS. The plates were incubated for 24 hr and then the inside of the chamber was gently wiped to remove the cells in the top chamber. The chamber was fixed with 4% PFS for 20 min, then incubated in 0.5% crystal violet in 20% methanol for 20 min with gentle shaking, and then washed with water. The chambers were air dried and the membranes were photographed. Using the microscope, representative photos were taken of each chamber. The number of migrated cells was counted manually in each photograph. Three independent experiments were performed with independent biologic replicates. Ordinary one-way ANOVA tests were used to evaluate for significant differences between different treatments.

### Preparation of whole cell lysates

OVCAR4 and OVCAR3 cells were cultured as described above and used to prepare whole cell lysates. Cells were washed twice with PBS solution and then resuspended in Lysis buffer (20 mM Tris-HCl pH 7.5, 150 mM NaCl, 1 mM EDTA, 1 mM EGTA, 1% NP-40, 1% sodium deoxycholate, 0.1% SDS). The cells were vortexed for 30 seconds and centrifuged for 10 were measured using the Bio-Rad Protein Assay Dye Reagent and volumes of lysates containing equal total amounts of protein were mixed with 4X SDS loading dye with Bromophenol blue. The lysates were then incubated for 5 min at 100°C.

### Western blotting

Equal amounts of whole cell lysates were prepared for SDS-PAGE gel as above and run on 8% SDS-PAGE gels. The gels were then transferred to nitrocellulose membranes and blocked with 5% nonfat milk in Tris-Buffered Saline with 0.1% Tween 20 (TBST) for 1 hr at room temperature. Primary antibodies were diluted in TBST and incubated for 1 hr at room temperature or at 4°C overnight. The membranes were washed multiple times in TBST and then incubated with horseradish peroxidase (HRP)-conjugated secondary antibody diluted in 5% nonfat milk in TBST for 1 hr at room temperature. Signals were captured using a luminol-based enhanced chemiluminescence HRP substrate (SuperSignal™ West Pico, Thermo Scientific) or an ultra-sensitive enhanced chemiluminescence HRP substrate (SuperSignal™ West Femto, Thermo Scientific) and a ChemiDoc imaging system (Bio-Rad).

### Immunoprecipitation and Western blotting of whole cell lysates

Immunoprecipitation of whole cell lysates was performed after plating OVCAR4 and OVCAR3 cells on 15 cm diameter cell culture dishes. Cells were treated with media supplemented with various concentrations of RBN-2397 and paclitaxel for 24 hr. Two 15 cm dishes were used for each treatment concentration. The cells were then washed with cold PBS, collected, and resuspended in IP Lysis Buffer (50 mM Tris-HCl pH 7.5, 0.5 M NaCl, 1.0 mM EDTA, 1% NP-40 and 10% glycerol, freshly supplemented with 1X phosphatase inhibitor cocktail 2, 1X phosphatase inhibitor cocktail 3, 1X complete protease inhibitor cocktail, 200 mM PMSF, 10 µM PJ34, 250 nM ADP-HDP, and 1mM DTT). The cells were vortexed for 30 seconds and centrifuged for 10 min at 4°C to remove the cellular debris, and the extracts were collected. The protein concentrations were then measured using Bradford assays. A small percentage of each cell lysate was saved for input. The cell lysates for IP were incubated with 3 µg of MAR detection reagent or rabbit IgG and protein A agarose beads overnight at 4°C with gentle rotating. The beads were then washed three times with lysis buffer for 5 min each at 4°C. The beads were then heated to 100°C for 5 min in 1X SDS-PAGE loading buffer to release the bound proteins. The immunoprecipitated material was then used for Western blotting.

### Generation of lentiviral expression vectors for wild-type and MARylation site mutant α-tubulin

The wild-type and MARylation site mutated α-tubulin cDNA was amplified from pCDNA3 clones as previously described [12] using primers encoding an N-terminal FLAG epitope tag listed below and cloned into the pINDUCER20 lentiviral doxycycline (Dox)-inducible expression vector (Addgene, 44012; RRID: Addgene_44012).

*pInducer-* α*-tubulin-F: 5’-TCCGCGGCCCCGAACTAGTGATGCGTGAGTGTATCTCTATC-3’*

*pInducer-* α*-tubulin-R: 5’-GTTTAATTAATCATTACTACTTAGGCGTAGTCAGGCACGTC-3’*

### Generation of cell lines with inducible ectopic expression of wild-type or MARylation site mutant **α**-tubulin

Cells were transduced with lentiviruses for Dox-inducible ectopic expression of α-tubulin (wild-type or mutant). We generated lentiviruses by transfection of the pINDUCER20 constructs described above, together with: (i) an expression vector for the VSV-G envelope protein (pCMV-VSV-G, Addgene 8454; RRID: Addgene_8454), (ii) an expression vector for GAG-Pol-Rev (psPAX2, Addgene 12260; RRID: Addgene_12260), and (iii) a vector to aid with translation initiation (pAdVAntage, Promega E1711) into HEK 293T cells using Lipofectamine 3000 reagent (Invitrogen, L3000015) according to the manufacturer’s instructions. The resulting viruses were collected in the culture medium, concentrated using Lenti-X Concentrator (Clontech, 631231), and used to infect OVCAR4 cells. Stably transduced cells were selected with G418 sulfate (Sigma, A1720; 1 mg/ml). The cells were treated with 1 µg/mL Dox for 24 hr to induce protein expression. Inducible ectopic expression of the cognate proteins was confirmed by Western blotting.

### Cell migration assays using OVCAR4 cells with inducible **α**-tubulin expression

Boyden chamber assays were used to determine the effects of various treatments on cell migration. Cells were treated with 1 µg/mL Dox for 24 hr to induce protein expression and manufacturer’s protocols. Cells were trypsinized, resuspended in serum-free media, and collected by centrifugation at 300 g for 3 min at room temperature. The cell pellets were then resuspended in serum-free media and 500 µL of media containing 100,000 cells were plated into the top chamber along with the indicated treatments. The chambers were then incubated in serum-containing media with FBS. The plates were incubated for 24 hr and then the inside of the chamber was gently wiped to remove the cells in the top chamber. The chamber was incubated in 0.5% crystal violet in 20% methanol for 30 min with gentle shaking and then washed with water. The chambers were air dried, and the membranes were photographed. Photographs were taken using a light microscope and the number of migrated cells were counted manually.

### Immunofluorescent staining

OVCAR4 and OVCAR3 cells were maintained in cell culture as described above. Cells were seeded at 4 x 10^4^ cells per well on 4-or 8-well chambered slides (ThermoFisher Scientific, 154534) one day prior to treatment. For Dox-induction, cells were treated with 1µg/mL Dox and incubated for 24 hr at 37°C. The cells were then treated with DMSO, RBN-2397 (5 µM), paclitaxel (1 nM), or combination. For cold treatment, cells were placed on ice for 45 min, washed with warm medium three times and allowed to recover in normal growth medium at 37°C for 15 min. For nocodazole treatment, nocodazole (MedChemExpress, HY-13520) was added at 6 µM for 1 hr. Cells were washed twice with PBS and fixed with 4% paraformaldehyde for 15 min at room temperature. Cells were then washed with PBS again and incubated at room temperature in Blocking Solution (PBS with 1% BSA, 0.3 M glycine, and 0.1% Tween-20) for 1 hr. The fixed cells were incubated with α-tubulin antibody or HA antibody in PBS overnight at 4°C. Cells were washed three times with PBS and then incubated with Alexa Fluor 594 goat anti-mouse IgG (1:500). The images were acquired using a confocal microscope and the fluorescence intensities were measured using Fiji Image J software. One percent outliers were removed, and ordinary one-way ANOVA tests were used to evaluate for significant differences between different treatments.

### RNA sequencing and data analysis

#### Generation of RNA-seq libraries

Total RNA was isolated from OVCAR4 and OCVAR3 the RNeasy kit (Qiagen, 74106) according to the manufacturer’s instructions. The NEBNext Ultra II Directional RNA Library Prep (NEB, E7765L) in conjunction with Poly(A) mRNA Magnetic Isolation Module kit (NEB, E7490L) was then used to prepare libraries according to the manufacturer’s instructions. The RNA-seq libraries were subjected to QC analyses [final library yields as assessed by Qubit 2.0 (Invitrogen) and size distribution of the final library DNA fragments as assessed by TapeStation (Illumina)] and sequenced using an Illumina HiSeq 2000.

#### Analysis of RNA-seq data

FASTQ files were subjected to quality control analyses using the FastQC tool. The reads were mapped to the human genome (hg38) using the STAR aligner, version 2.7.2b [14]. Transcriptome assembly was performed using cufflinks v.2.2.1 with default parameters [15]. The transcripts were merged into distinct, non-overlapping set using cuffmerge, followed by cuffdiff to call the differentially regulated transcripts [16]. The significantly regulated genes upon RBN-2397, paclitaxel, or combination treatment compared to vehicle were associated with a p-value less than 0.05.

#### Transcriptome data analyses

A fold change cutoff of 1.5 was used for upregulated genes and fold change of 0.67 and lower were termed downregulated genes in all treatments. Biovenn was used to construct Venn diagrams for up and down regulated genes grouped together in each treatment arm.

#### Gene Ontology (GO) analyses

Gene ontology analyses were conducted using the Database for Annotation, Visualization, and Integrated Discovery (DAVID) Bioinformatics Resources tool [17, 18] for genes specifically enriched in combination treatment. Ontological terms are ranked based on enrichment score.

### Data availability statement

The data in this study are available upon request from the corresponding author. RNA-seq data has been published and can be accessed from GEO using accession number GSE268985.

## Results

### RBN-2397 in combination with paclitaxel has a robust impact on ovarian cancer cell growth

Our previously published work demonstrated that PARP7 knockdown led to the reduction of ovarian cancer cell growth [12]. We sought to investigate the effect of PARP7 inhibition on ovarian cancer cell proliferation. We also wanted to determine the effect of paclitaxel on cancer cell growth. A dose response assay, where we treated the ovarian cancer cells (OVCAR4 and OVCAR3) with increasing doses of RBN-2397 or paclitaxel, showed a respective increase in cell death (Figure 1A and 1B). Considering the resistance to chemotherapy and the toxic effects of paclitaxel, along with the selectivity of RBN-2397 for PARP7, we selected doses for both paclitaxel and RBN-2397 that are below their respective IC50 thresholds for the growth assays. Single treatments of 500 nM RBN-2397 or 0.4 nM paclitaxel caused a significant decrease in proliferation of OVCAR4 cells but not OVCAR3 cells. Interestingly, the combined administration of both treatments resulted in a more pronounced decrease in proliferation for both OVCAR4 and OVCAR3 cell lines (Figure 1C and 1D). Our data suggests that RBN-2397 and paclitaxel combination treatment can more effectively decrease the growth of ovarian cancer cell lines.

**Figure 1.**
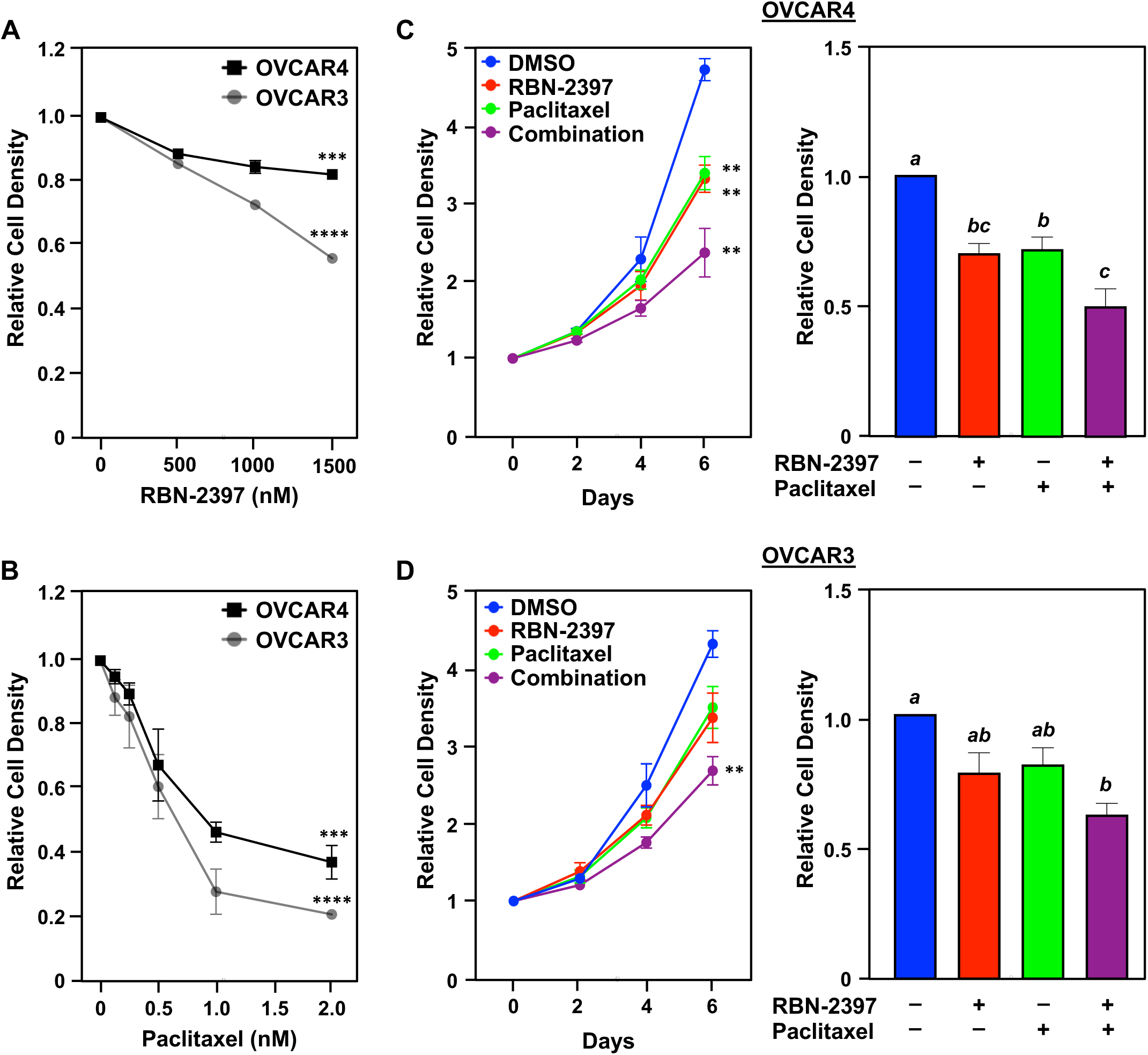
Impact of RBN-2397, paclitaxel, and their combination on ovarian cancer cell growth. **(A and B)** Dose response curves showing increasing concentrations of RBN-2397 (A) or paclitaxel (B) significantly decreasing cell viability in ovarian cancer cells (OVCAR4 and OVCAR3), assayed by crystal violet staining. IC50s were calculated; Paclitaxel: 0.47 nM (OVCAR4), 0.46 nM (OVCAR3); RBN-2397: 727.2 nM (OVCAR4), 1159 nM (OVCAR3). Points marked with asterisks are significantly different; Student’s t-test; *** = p<0.001, **** = p<0.0001. **(C and D)** Line graphs showing the growth of OVCAR4 cells (C) and OVCAR3 cells (D) over a period of 6 days with different conditions: DMSO, RBN-2397 (500 nM), paclitaxel (0.4 nM), combination (*left panels*). Each point represents the mean ± SEM; n=3. Points marked with asterisks are significantly different; Student’s t-test; ** = p<0.01. Bar graphs showing quantification of the proliferation assays at Day 6 (*right panels*). Each bar represents the mean ± SEM; n=3. Bars marked with different letters are significantly different, Ordinary one-way ANOVA test.

### The migratory phenotype of ovarian cancer cells is nearly abolished by combining RBN-2397 and paclitaxel

PARP7 depletion has also been shown to reduce OVCAR4 and OVCAR3 cell migration [12]. To evaluate whether PARP7 inhibition by RBN-2397 has the same phenotypic effect, we performed cell migration assays. OVCAR4 and OVCAR3 cells were plated in Boyden chamber assays as previously described and treated with RBN-2397, paclitaxel, or combination. RBN-2397 treatment alone resulted in a significant decrease in migration of OVCAR3 cells but not in OVCAR4 cells (Figure 2A and 2B; Figure S1A and S1B). Paclitaxel single treatment significantly reduced migration of both OVCAR4 and OVCAR3 cells. Remarkably, the combination treatment led to a more significant decrease in the migration of ovarian cancer cells than either of the treatments alone (Figure 2A and 2B; Figure S1A and S1B). This data supports a role for PARP7 enzymatic activity and microtubule assembly in ovarian cancer cell migration.

**Figure 2.**
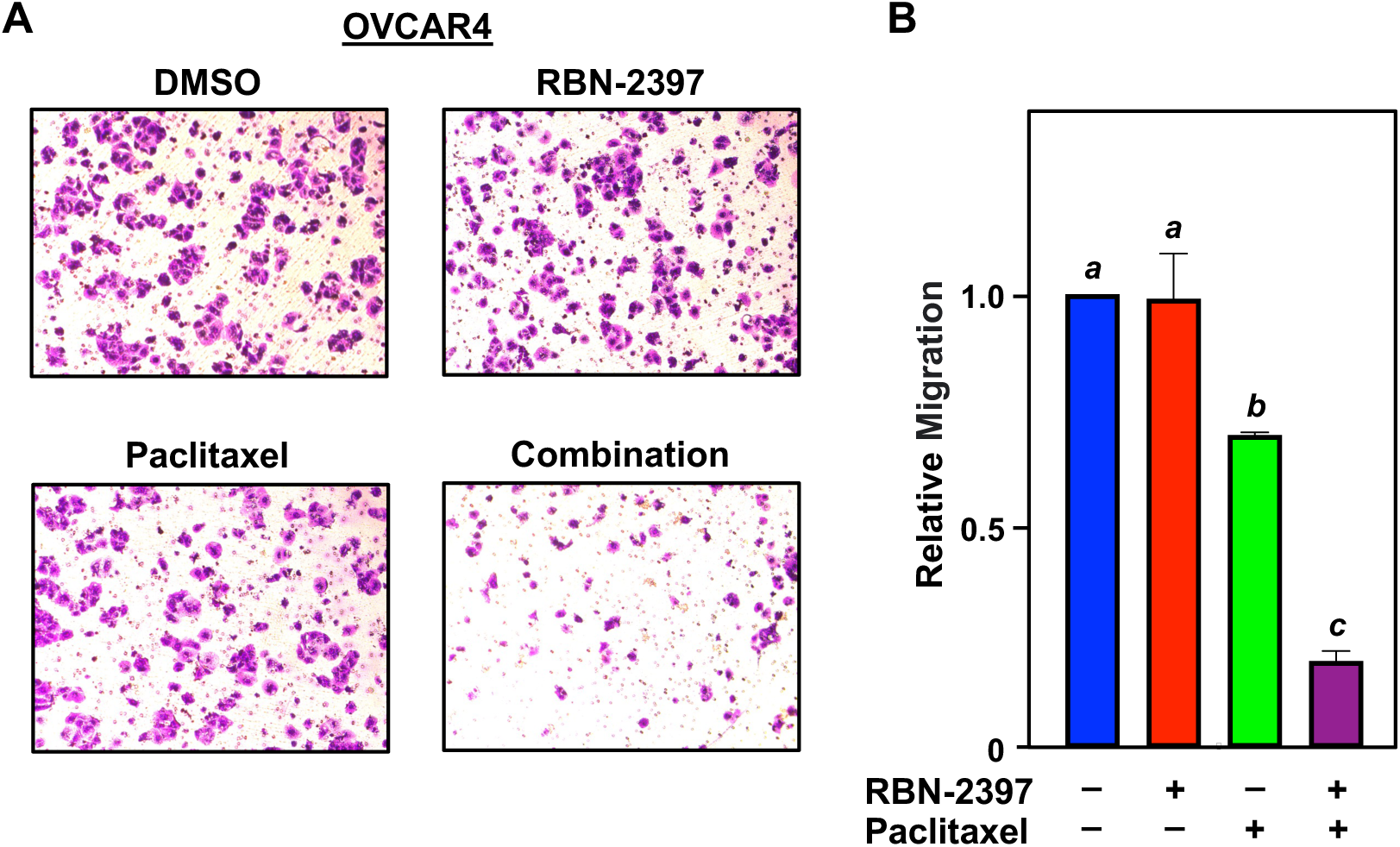
Effect of RBN-2397, paclitaxel, and their combination on ovarian cancer cell migration. **(A)** Representative images of cell migration assays for OVCAR4 cells treated with DMSO, RBN-2397 (500 nM), paclitaxel (0.4 nM), and combination. **(B)** Quantification of the cell migration assays shown in (A) for OVCAR4 cells. Each bar represents the mean ± SEM; n=3. Bars marked with different letters are significantly different, Ordinary one-way ANOVA test.

### RBN-2397 blocks the MARylation of **α**-tubulin and stabilizes the tubulin network

One known target of PARP7 is autoMARylation. To evaluate whether RBN-2397 directly inhibits the effects of PARP7, we examined the effect of RBN-2397 on PARP7 autoMARylation. Using a FLAG-tagged PARP7 cDNA, PARP7 was ectopically expressed in OVCAR4 and OVCAR3 cells, then treated with vehicle or RBN-2397. Immunoprecipitation of FLAG-tagged PARP7 and immunoblotting for MAR showed that the RBN-2397 significantly blocks MARylation of PARP7 (Figure 3A; Figure S2A). In our previous studies, we utilized a recombinant analog-sensitive PARP7 and mass spectrometry approach to reveal that α-tubulin is subject to PARP7-mediated MARylation [12]. We were interested in evaluating whether PARP7 inhibition by RBN-2397 would block this activity. We treated ovarian cancer cells with RBN-2397 and evaluated α-tubulin MARylation. At endogenous levels of PARP7, RBN-2397 (alone or in combination) significantly blocked the MARylation of α-tubulin. Interestingly, we observed an increase in α-tubulin MARylation upon treatment with paclitaxel compared to control. (Figure 3B). Hence, we wanted to investigate whether RBN-2397 and/or paclitaxel influence PARP7 protein levels. Although not statistically significant, we observed that RBN-2397 and paclitaxel stabilized PARP7 protein (Figure 3C and 3D; Figure S2B and S2C). These results demonstrate that PARP7 inhibition blocks α-tubulin MARylation even in the presence of paclitaxel.

**Figure 3.**
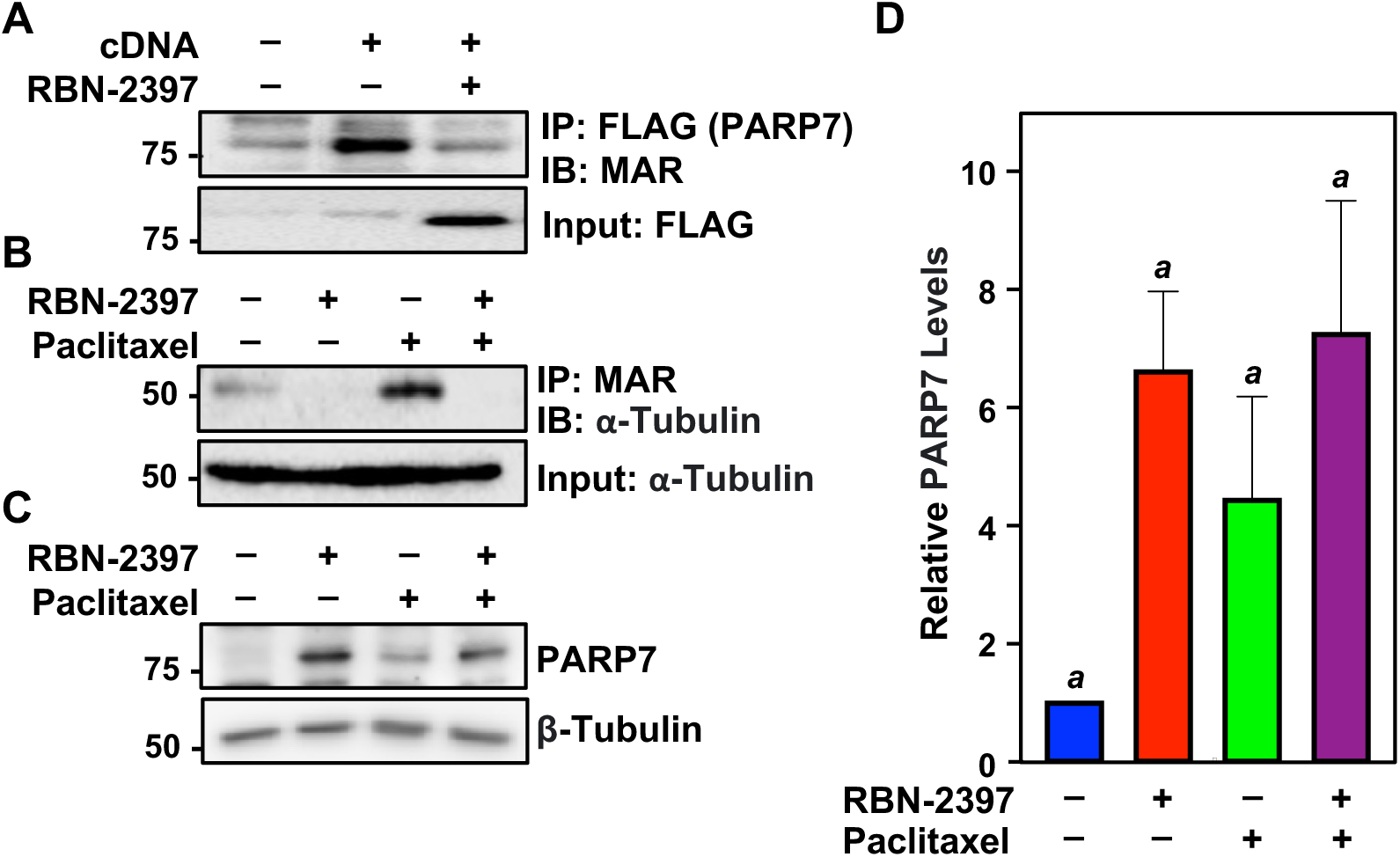
RBN-2397 blocks PARP7 autoMARylation and the MARylation of α-tubulin. **(A)** RBN-2397 blocks the autoMARylation of ectopically expressed FLAG-PARP7 in OVCAR4 cells, assayed by immunoprecipitation of FLAG followed by Western blot for MAR. **(B)** RBN-2397 blocks the MARylation of α-tubulin in OVCAR4 cells, even in the presence of paclitaxel, assayed by immunoprecipitation of MAR followed by Western blot for α-tubulin. **(C)** Stabilization of endogenous PARP7 levels were detected in OVCAR4 cells following treatment with RBN-2397 (1 μM) alone and in combination with paclitaxel (1 nM), assayed by Western blot. **(D)** Bar graph showing quantification of PARP7 levels from experiments shown in panel (C) for OVCAR4 cells. Each bar represents the mean ± SEM; n=3. Bars marked with different letters are significantly different, Ordinary one-way ANOVA test.

Given that RBN-2397 and paclitaxel affect microtubule activities, and that microtubules have key roles in cellular migration, we aimed to evaluate their effect as single agents and in combination on α-tubulin stability in ovarian cancer cells. To assess this, we examined the recovery from microtubule depolymerization in OVCAR4 and OVCAR3 cells treated with vehicle, RBN-2397, paclitaxel, or combination. We conducted immunostaining for α-tubulin and visualized the cells using fluorescence microscopy (Figure 4A; Figure S3A). After initial exposure to cold temperature to destabilize the microtubules, the cells were shifted to 37°C to facilitate microtubule regrowth. RBN-2397 and paclitaxel single treatments led to the stabilization of the cold-destabilized microtubule structures containing α-tubulin. Comparable results were observed in a parallel experiment involving cells treated with nocodazole, a drug known to depolymerize microtubules. Notably, we observed a more significant increase in α-tubulin stability with RBN-2397 and paclitaxel combination treatment after cold or nocodazole exposure (Figure 4A and 4B; Figure S3A and S3B). Collectively, our data suggests that inhibiting the catalytic activity of PARP7 by RBN-2397 blocks PARP7-mediated α-tubulin MARylation, which in turn stabilizes the microtubule network. Interestingly, the addition of paclitaxel further stabilizes the network potentially explaining the cooperative effect of RBN-2397 and paclitaxel seen on ovarian cancer cell migration.

**Figure 4.**
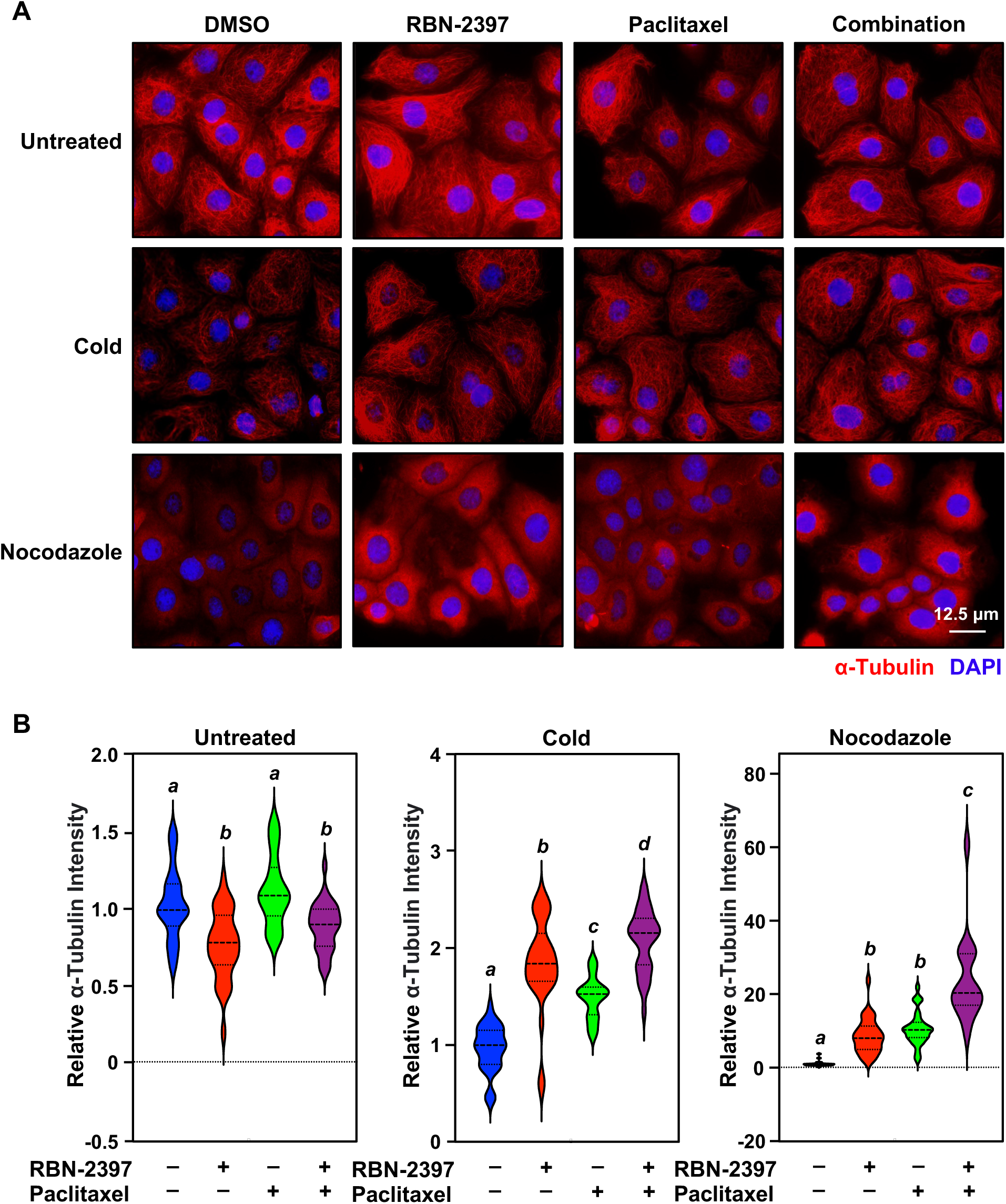
Effect of RBN-2397, paclitaxel, and their combination on α-tubulin network after inducing microtubule destabilization. **(A)** RBN-2397, paclitaxel, and their combination promote microtubule stability in OVCAR4 cells. Representative immunofluorescent images of α-tubulin in OVCAR4 cells after treatment with DMSO, RBN-2397 (500 nM), paclitaxel (0.4 nM), or their combination and exposure to cold treatment or nocodazole. The experiment was performed 3 times to ensure reproducibility. Scale bar = 12.5 μm. **(B)** Violin plots quantifying α-tubulin staining from experiments shown in (A) in OVCAR4 cells untreated *(left panel)*, cold *(middle panel)*, or nocodazole treated *(right panel)*. Violins marked with different letters are significantly different, Ordinary one-way ANOVA test; n=3.

### Mutating the MARylation site in **α**-tubulin stabilizes the tubulin network and blocks ovarian cancer cell migration in the presence of paclitaxel

Prior work has attributed the decrease in cell migration upon PARP7 knockdown to stabilization of the tubulin network in the absence of PARP7, restricting growth and migration. To confirm that inhibition of α-tubulin MARylation is directly responsible for increasing tubulin stability, we engineered OVCAR4 cell lines expressing wild-type or site-specific mutant α-tubulin to test whether mutating the MARylation site would have the same effect as RBN-2397 (Figure 5A). After inducing ectopic expression of wild-type or mutant α-tubulin, cells were treated with vehicle or paclitaxel, and immunofluorescence assays were used to measure the levels of α-tubulin. α-tubulin was, in fact, stabilized with mutation of the MARylation site after cold exposure. The addition of paclitaxel further stabilized the microtubule network (Figure 5B and 5C), corroborating our observations with RBN-2397.

**Figure 5.**
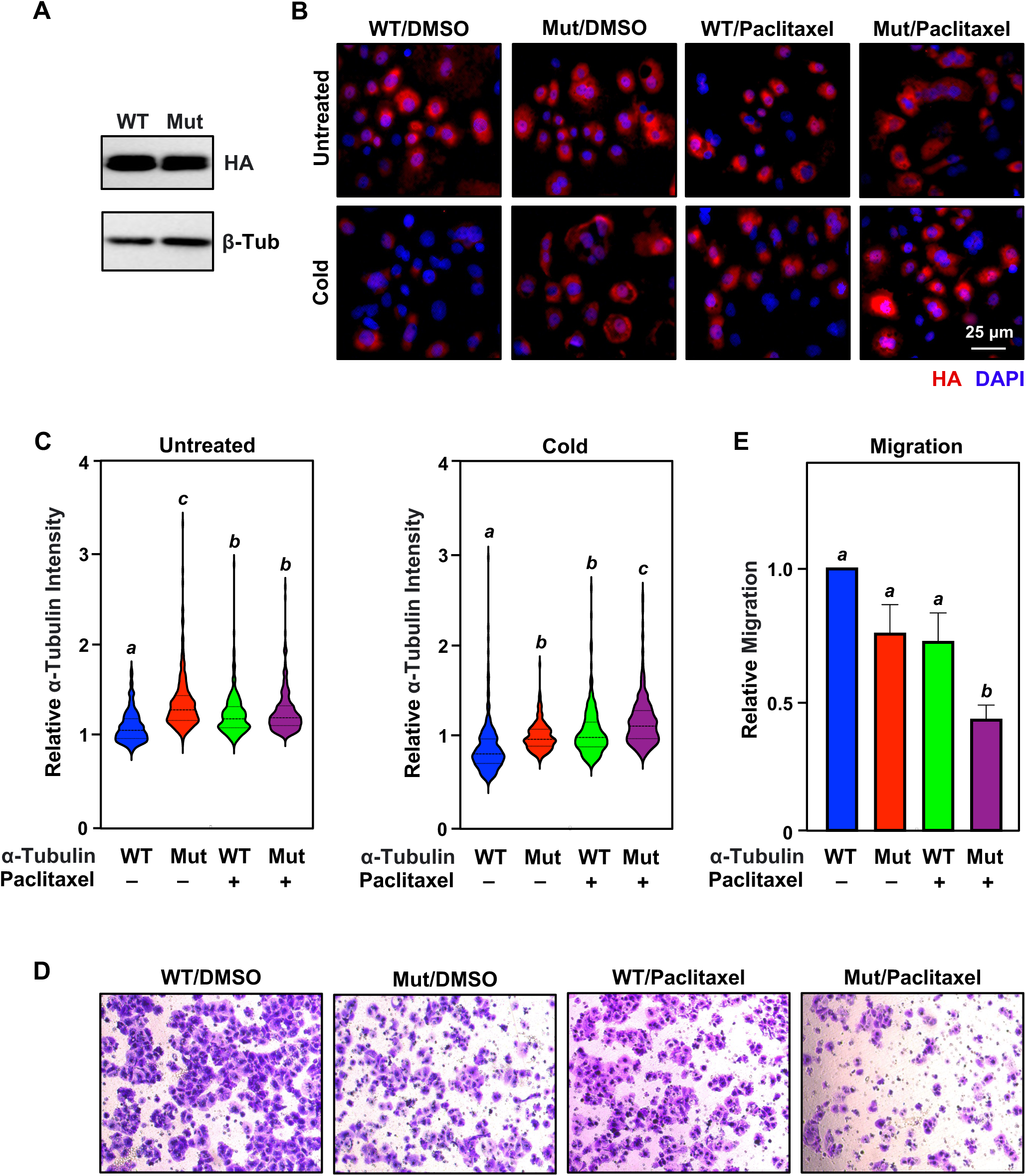
Mutating the MARylation site in α-tubulin synergizes with paclitaxel to stabilize the tubulin network and block ovarian cancer cell migration. **(A)** Wild-type (WT) and MARylation-site mutant α-tubulin (Mut) are expressed at similar levels in the wild-type and mutant OVCAR4 cell lines, assayed by Western blot. **(B)** Representative immunofluorescent images of α-tubulin showing increased microtubule stability in Mut compared to WT; paclitaxel (1 nM) enhances microtubule stability in both mutant and WT after cold exposure. The experiment was performed 3 times to ensure reproducibility. Scale bar = 25 μm. **(C)** Violin plots quantifying α-tubulin staining from experiments shown in (B) in untreated *(left panel)* and cold treated cells *(right panel)*. Violins marked with different letters are significantly different, Ordinary one-way ANOVA test; n=3. **(D and E)** Cell migration assays for OVCAR4 WT and α-tubulin mutant with DMSO and paclitaxel (1 nM). Bar plot showing quantification of the cell migration assays shown in (D). Each bar represents the mean ± SEM; n=3. Bars marked with different letters are significantly different, Ordinary one-way ANOVA test.

We hypothesized that mutating the MARylation site on α-tubulin would produce the same phenotypic changes in cell migration as PARP7 inhibition. Hence, OVCAR4 cell lines expressing wild-type or mutant α-tubulin were plated in Boyden chamber assays as previously described and treated with paclitaxel. OVCAR4 cells expressing the α-tubulin mutant showed significantly decreased migration in the presence of paclitaxel compared to wild-type, mimicking the phenotype observed with PARP7 inhibition and paclitaxel (Figure 5D and 5E). This supports the role of site-specific MARylation of α-tubulin by PARP7 in ovarian cancer cell migration.

### Effect of RBN-2397, paclitaxel, and their combination on ovarian cancer cell biology

PARP7 has been implicated in the suppression of type I interferon response – and therefore T cell mediated immunity – as well as ovarian cancer cell growth and migration [9]. Our work above suggests a robust response in ovarian cancer cells to PARP7 inhibition in combination with paclitaxel. RNA-sequencing data of siRNA-mediated PARP7 knockdown in OVCAR4 human ovarian cancer cells from our previous study revealed an enrichment of genes encoding proteins involved in cell-cell adhesion, cell cycle arrest, apoptosis, and gene regulation [12]. To further validate, we performed RNA-sequencing in both OVCAR4 and OVCAR3 cells treated with RBN-2397, paclitaxel, or combination. Although the number of genes that were differentially regulated was low (Figure S4A and S4C), evaluating the ontologies of the genes that were up-or down regulated by the combination treatment yielded terms related to cellular proliferation, cell-cell adhesion, and chemotaxis reinforcing our previous observations (Figure S4B and S4D). Altogether, we propose that the mechanism of synergy between RBN-2397 and paclitaxel is a cytoplasmic process involving the microtubules and does not directly involve gene regulation.

## Discussion

Our results, in this study, highlight the role of PARP7-mediated ADP-ribosylation in ovarian cancer cell biology. We have previously shown that PARP7 knockdown decreases ovarian cancer cell proliferation and migration by stabilizing α-tubulin. Herein, we show that inhibiting the catalytic activity of PARP7 by RBN-2397 mimics the effect of PARP7 depletion and stabilizes α-tubulin to inhibit ovarian cancer cell proliferation and migration. RBN-2397 synergizes with paclitaxel to further stabilize the microtubules and inhibit the aggressive phenotypes of high-grade ovarian cancer cells. Collectively, we propose that the synergistic effect between RBN-2397 and paclitaxel indicates a potential therapeutic target for high grade ovarian cancers.

### Current treatments and challenges for recurrent ovarian cancer

Treatment options for recurrent ovarian cancer are limited, with low response rates, limited response duration, and significant toxicity profiles [19]. More effective, targeted treatments are needed, particularly in the platinum resistant setting. The most effective systemic treatment for platinum-resistant ovarian cancer remains paclitaxel, although many tumors are or become resistant to taxanes [19]. As patients receiving treatment for recurrent ovarian cancer are often facing cumulative toxicities and worsening cancer burden, a low effective dose of paclitaxel in combination with other agents is desirable. Our study provides evidence that PARP7, which plays a role in MARylation and destabilization of α-tubulin, can be targeted with RBN-2397, a PARP7 inhibitor, to improve sensitivity to paclitaxel in platinum resistant ovarian cancer cell lines in vitro. Notably, the work in this study was done using very low doses of paclitaxel.

### Role of PARP7 in ovarian cancer biology

While some members of the PARP family of enzymes play clear roles in cancer biology and have been identified for targeted treatments in ovarian cancer, a full understanding of the role of these enzymes is still being elucidated. PARP7, a mono (ADP-ribosyl) transferase, has been implicated in cancer biology. Prior work has shown that PARP7 MARylates α-tubulin, promoting microtubule instability [12]. This led us to evaluate whether the PARP7 inhibitor, further examined the effect of RBN-2397 in combination with paclitaxel, which also affects microtubules.

Both catalytic inhibition of PARP7 and a MARylation site-specific mutant of α-tubulin resulted in significantly decreased ovarian cancer cell migration when combined with paclitaxel. We also explored changes in cellular biology following these treatments alone or in combination by RNA sequencing. The number of genes that were differentially regulated was low, but GO analysis revealed that RBN-2397 and paclitaxel combination treatment enriched terms related to cellular proliferation, cell-cell adhesion, and chemotaxis reinforcing our previous observations.

Although RBN-2397 is a selective inhibitor of PARP7 that blocks its enzymatic activity, we found that PARP7 inhibition resulted in increased levels of PARP7 protein. PARP7 has been shown to be a relatively unstable protein, with rapid turnover compared to other PARP family members [20]. All three domains – the zinc finger, tryptophan-tryptophan-glutamate (WWE) domain, and catalytic domain – contribute to the short half-life of the protein [20]. Notably, introducing loss-of-function mutations into the PARP7 catalytic domain has been shown to increase PARP7 protein levels in both breast cancer and prostate cancer cell lines [20, 21]. Kamata *et. al.* found that mutations in the catalytic domain increased the half-life of PARP7 approximately 8-fold [20]. In our work, PARP7 inhibition also increased PARP7 protein levels. This increase in PARP7 levels with PARP7 inhibition is not unique to ovarian cancer cells, as it has also been observed in breast cancer cells treated with RBN-2397 [21]. Given these findings, it is likely that PARP7 activity itself, which includes autoMARylation, is necessary for PARP7 degradation.

### Mechanistic insights and future directions

PARP7 knockdown has been shown to significantly impact both cell proliferation and migration [12]. PARP7 inhibition by RBN-2397 shows a similar impact on ovarian cancer cell proliferation and this effect was intensified by the addition of paclitaxel. The effect of PARP7 inhibition on migration was significant in OVCAR3 cell line. The addition of paclitaxel resulted in a striking decrease in ovarian cancer cell migration, in both OVCAR4 and OVCAR3. RNA sequencing showed genes related to cellular proliferation, cell-cell adhesion, and chemotaxis were up-or downregulated with RBN-2397 and paclitaxel combination treatment. Previously, baseline microtubule stability has been shown to correlate with paclitaxel-mediated microtubule stabilization and ovarian cancer cell response to paclitaxel [22]. Paclitaxel-resistant cell lines isolated from SKOV3 and OVCAR3 have been shown to have enhanced microtubule dynamics and decreased polymerized microtubules compared to the parent, non-paclitaxel resistant lines [23]. Depletion of multiple kinases, which resulted in microtubule stabilization in vitro, has been shown to similarly enhance paclitaxel sensitivity in vitro and in human ovarian cancer xenograft models [24]. Mechanistically, the impact of PARP7 inhibition on ovarian cancer cell migration may be explained by the stabilization of α-tubulin. It is possible that microtubule stability enhances sensitivity to paclitaxel by increasing the microtubule framework for taxane binding. The role of microtubule stability in increasing paclitaxel sensitivity can be leveraged to improve response to paclitaxel in ovarian cancer cells.

In conclusion, the link between α-tubulin MARylation and stabilization and increased paclitaxel sensitivity provides a potential therapeutic target for platinum resistant ovarian cancer, a disease with few effective treatment options. Further studies of PARP7 inhibition in combination with other microtubule-stabilizing and destabilizing agents, such as docetaxel or Vinca alkaloids, are also warranted. Moving forward, to build upon these promising findings in vitro, we hope to validate the combination of PARP7 inhibition and paclitaxel on tumor growth using *in vivo* models prior to phase I clinical studies.

## Authors’ Contributions

A.N.S. – conceptualization, methodology, formal analysis, investigation, writing-original draft, writing-review and editing, visualization; M.W.A. – methodology, formal analysis, investigation, writing-original draft, writing-review and editing, visualization; S.C. – methodology, supervision; S.K. – formal analysis; J.S.L. – Resources; W.L.K. – conceptualization, validation, writing-review and editing, supervision, project administration, funding acquisition; C.V.C. – methodology, validation, writing-review and editing, supervision, project administration.

## Acknowledgments

We would like to acknowledge the UT Southwestern Next Generation Sequencing Core, under the direction of Ralf Kittler. This work was supported by grants from the NIH/National Institute of Diabetes and Digestive and Kidney Diseases (NIH/NIDDK; R01 DK069710 to W.L.K.), the U.S. Department of Defense (DOD) Ovarian Cancer Research Program (OCRP; OC200311 and OC230196 to W.L.K.), and funds from the Cecil H. and Ida Green Center for Reproductive Biology Sciences Endowment to W.L.K.

## Note

Supplementary Materials for this article are available.

## References

1. Siegel RL, Giaquinto AN, Jemal A (2024) Cancer statistics, 2024. CA Cancer J Clin 74:12-49.

2. Burke W, Barkley J, Barrows E, Brooks R, Gecsi K, Huber-Keener K, Jeudy M, Mei S, et al. (2023) Executive summary of the ovarian cancer evidence review conference. Obstetrics and Gynecology 142:179.

3. Poveda AM, Selle F, Hilpert F, Reuss A, Savarese A, Vergote I, Witteveen P, Bamias A, et al. (2015) Bevacizumab combined with weekly paclitaxel, pegylated liposomal doxorubicin, or topotecan in platinum-resistant recurrent ovarian cancer: analysis by chemotherapy cohort of the randomized phase III AURELIA trial. Journal of Clinical Oncology 33:3836–3838.

4. Ortiz M, Wabel E, Mitchell K, Horibata S (2022) Mechanisms of chemotherapy resistance in ovarian cancer. Cancer Drug Resistance 5:304.

5. Maloney SM, Hoover CA, Morejon-Lasso LV, Prosperi JR (2020) Mechanisms of taxane resistance. Cancers 12:3323.

6. Sanderson DJ, Cohen MS (2020) Mechanisms governing PARP expression, localization, and activity in cells. Critical Reviews in Biochemistry and Molecular Biology 55:541–554.

7. Challa S, Stokes MS, Kraus WL (2021) MARTs and MARylation in the cytosol: biological functions, mechanisms of action, and therapeutic potential. Cells 10:313.

8. Wu Y, Xu S, Cheng S, Yang J, Wang Y (2023) Clinical application of PARP inhibitors in ovarian cancer: from molecular mechanisms to the current status. Journal of Ovarian Research 16:6.

9. Gozgit JM, Vasbinder MM, Abo RP, Kunii K, Kuplast-Barr KG, Gui B, Lu AZ, Molina JR, et al. (2021) PARP7 negatively regulates the type I interferon response in cancer cells and its inhibition triggers antitumor immunity. Cancer Cell 39:1214–1226. e1210.

10. MacPherson L, Tamblyn L, Rajendra S, Bralha F, McPherson JP, Matthews J (2013) 2, 3, 7, 8-Tetrachlorodibenzo-p-dioxin poly (ADP-ribose) polymerase (TiPARP, ARTD14) is a mono-ADP-ribosyltransferase and repressor of aryl hydrocarbon receptor transactivation. Nucleic acids research 41:1604-1621.

11. Bindesbøll C, Tan S, Bott D, Cho T, Tamblyn L, MacPherson L, Grønning-Wang L, Nebb HI, et al. (2016) TCDD-inducible poly-ADP-ribose polymerase (TIPARP/PARP7) mono-ADP-ribosylates and co-activates liver X receptors. Biochemical Journal 473:899–910.

12. Palavalli Parsons LH, Challa S, Gibson BA, Nandu T, Stokes MS, Huang D, Lea JS, Kraus WL (2021) Identification of PARP-7 substrates reveals a role for MARylation in microtubule control in ovarian cancer cells. Elife 10:e60481.

13. Gibson BA, Conrad LB, Huang D, Kraus WL (2017) Generation and characterization of recombinant antibody-like ADP-ribose binding proteins. Biochemistry 56:6305–6316.

14. Dobin A, Davis CA, Schlesinger F, Drenkow J, Zaleski C, Jha S, Batut P, Chaisson M, et al. (2013) STAR: ultrafast universal RNA-seq aligner. Bioinformatics 29:15–21.

15. Trapnell C, Williams BA, Pertea G, Mortazavi A, Kwan G, Van Baren MJ, Salzberg SL, Wold BJ, et al. (2010) Transcript assembly and quantification by RNA-Seq reveals unannotated transcripts and isoform switching during cell differentiation. Nature biotechnology 28:511–515.

16. Trapnell C, Williams BA, Pertea G, Mortazavi A, Kwan G, van Baren MJ, Salzberg SL, Wold BJ, et al. (2010) Transcript assembly and abundance estimation from RNA-Seq reveals thousands of new transcripts and switching among isoforms. Nature biotechnology 28:511.

17. Sherman BT, Hao M, Qiu J, Jiao X, Baseler MW, Lane HC, Imamichi T, Chang W (2022) DAVID: a web server for functional enrichment analysis and functional annotation of gene lists (2021 update). Nucleic acids research 50:W216-W221.

18. Sherman BT, Lempicki RA (2009) Systematic and integrative analysis of large gene lists using DAVID bioinformatics resources.

19. Davis A, Tinker AV, Friedlander M (2014) “Platinum resistant” ovarian cancer: what is it, who to treat and how to measure benefit? Gynecologic oncology 133:624–631.

20. Kamata T, Yang C-S, Melhuish TA, Frierson Jr HF, Wotton D, Paschal BM (2021) Post-transcriptional regulation of PARP7 protein stability is controlled by androgen signaling. Cells 10:363.

21. Rasmussen M, Tan S, Somisetty VS, Hutin D, Olafsen NE, Moen A, Anonsen JH, Grant DM, et al. (2021) PARP7 and mono-ADP-ribosylation negatively regulate estrogen receptor α signaling in human breast cancer cells. Cells 10:623.

22. Ahmed AA, Wang X, Lu Z, Goldsmith J, Le X-F, Grandjean G, Bartholomeusz G, Broom B, et al. (2011) Modulating microtubule stability enhances the cytotoxic response of cancer cells to Paclitaxel. Cancer research 71:5806–5817.

23. McGrail DJ, Khambhati NN, Qi MX, Patel KS, Ravikumar N, Brandenburg CP, Dawson MR (2015) Alterations in ovarian cancer cell adhesion drive taxol resistance by increasing microtubule dynamics in a FAK-dependent manner. Scientific reports 5:9529.

24. Yang H, Mao W, Rodriguez-Aguayo C, Mangala LS, Bartholomeusz G, Iles LR, Jennings NB, Ahmed AA, et al. (2018) Paclitaxel sensitivity of ovarian cancer can be enhanced by knocking down pairs of kinases that regulate MAP4 phosphorylation and microtubule stability. Clinical Cancer Research 24:5072–5084.

