## Supplemental Materials for "RBN-2397, a PARP7 Inhibitor, Synergizes with Paclitaxel to Inhibit Proliferation and Migration of Ovarian Cancer Cells"

**Spirtos, Aljardali *et al.* (2024)**

This document contains the following supplemental information: Page

**Supplemental Figures** .......................................................2

- Figure S1. Related to Figure 2. Effect of RBN-2397, paclitaxel, and their combination on ovarian cancer cell migration. ..............................2
- Figure S2. Related to Figure 3. RBN-2397 blocks PARP7 auto-MARylation and the MARylation of α-tubulin. ..............................3
- Figure S3. Related to Figure 4. Effect of RBN-2397, paclitaxel, and their combination on α-tubulin network after inducing microtubule destabilization. ...........4
- Figure S4. Effect of RBN-2397, paclitaxel, and their combination on ovarian cancer cell biology..............................6

### Supplemental Figures

**Supplementary Figure 1. Effect of RBN-2397, paclitaxel, and their combination on ovarian cancer cell migration.**

**(A)** Representative images of cell migration assays for OVCAR3 cells treated with DMSO, RBN-2397 (500 nM), paclitaxel (0.4 nM), and combination.

**(B)** Quantification of the cell migration assays shown in (A) for OVCAR3 cells. Each bar represents the mean + SEM; n=3. Bars marked with different letters are significantly different, Ordinary one-way ANOVA test.

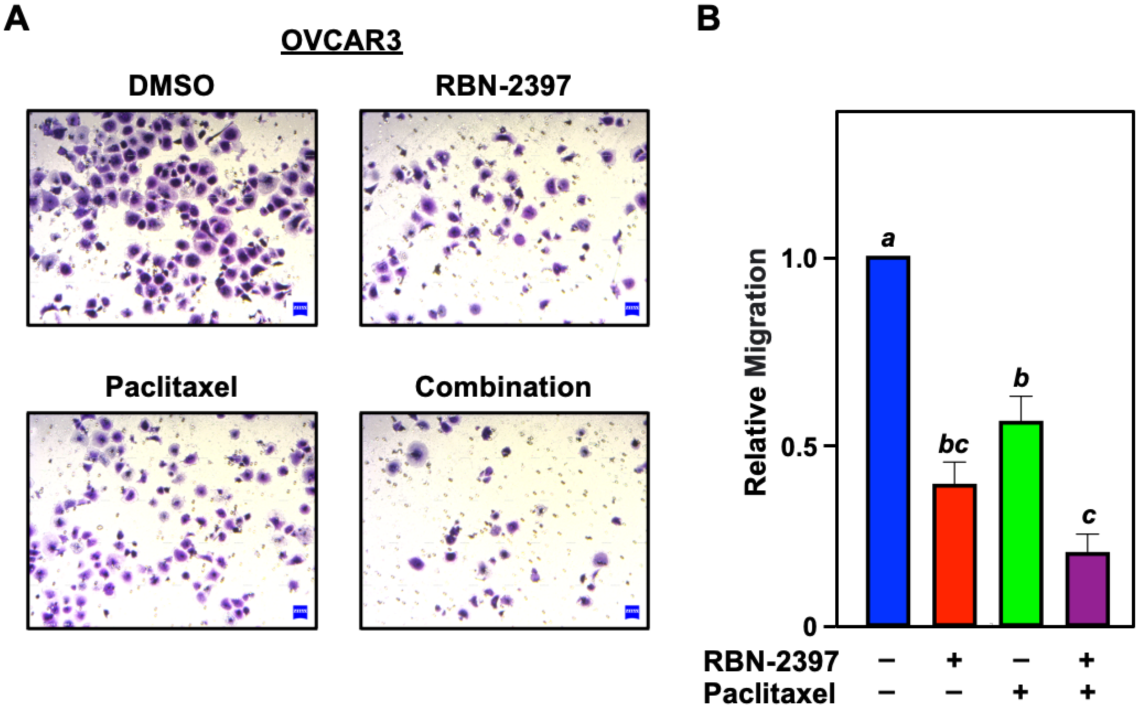

**Supplementary Figure 2. RBN-2397 blocks PARP7 auto-MARylation and the MARylation of α-tubulin.**

**(A)** RBN-2397, blocks the autoMARylation of ectopically expressed FLAG-PARP7 in OVCAR3 cells, assayed by immunoprecipitation of FLAG followed by Western blot for MAR.

**(B)** Stabilization of endogenous PARP7 levels were detected in OVCAR3 cells following treatment with RBN-2397 (1 μM) alone and in combination with paclitaxel (1 nM), assayed by Western blot.

**(C)** Bar graph showing quantification of PARP7 levels from experiments shown in panel (B) for OVCAR3 cells. Each bar represents the mean + SEM; n=3. Bars marked with different letters are significantly different, Ordinary one-way ANOVA test.

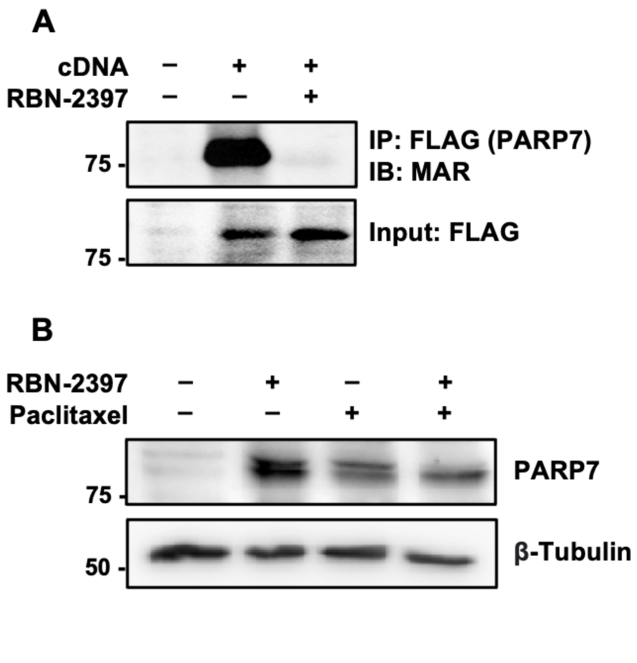

**Supplementary Figure 3. Effect of RBN-2397, paclitaxel, and their combination on α-tubulin network after inducing microtubule destabilization.**

**(A)** RBN-2397, paclitaxel and their combination promote microtubule stability in OVCAR3 cells. Representative immunofluorescent images of α-tubulin in OVCAR3 cells after treatment with DMSO, RBN-2397 (500 nM), paclitaxel (0.4 nM), or their combination and exposure to cold treatment or nocodazole. The experiment was performed 3 times to ensure reproducibility. Scale bar = 12.5 μm.

**(B)** Violin plots quantifying α-tubulin staining from experiments shown in (A) in OVCAR3 cells; untreated *(left panel)*, cold *(middle panel)*, or nocodazole treated *(right panel)*. Violins marked with different letters are significantly different, Ordinary one-way ANOVA test; n=3.

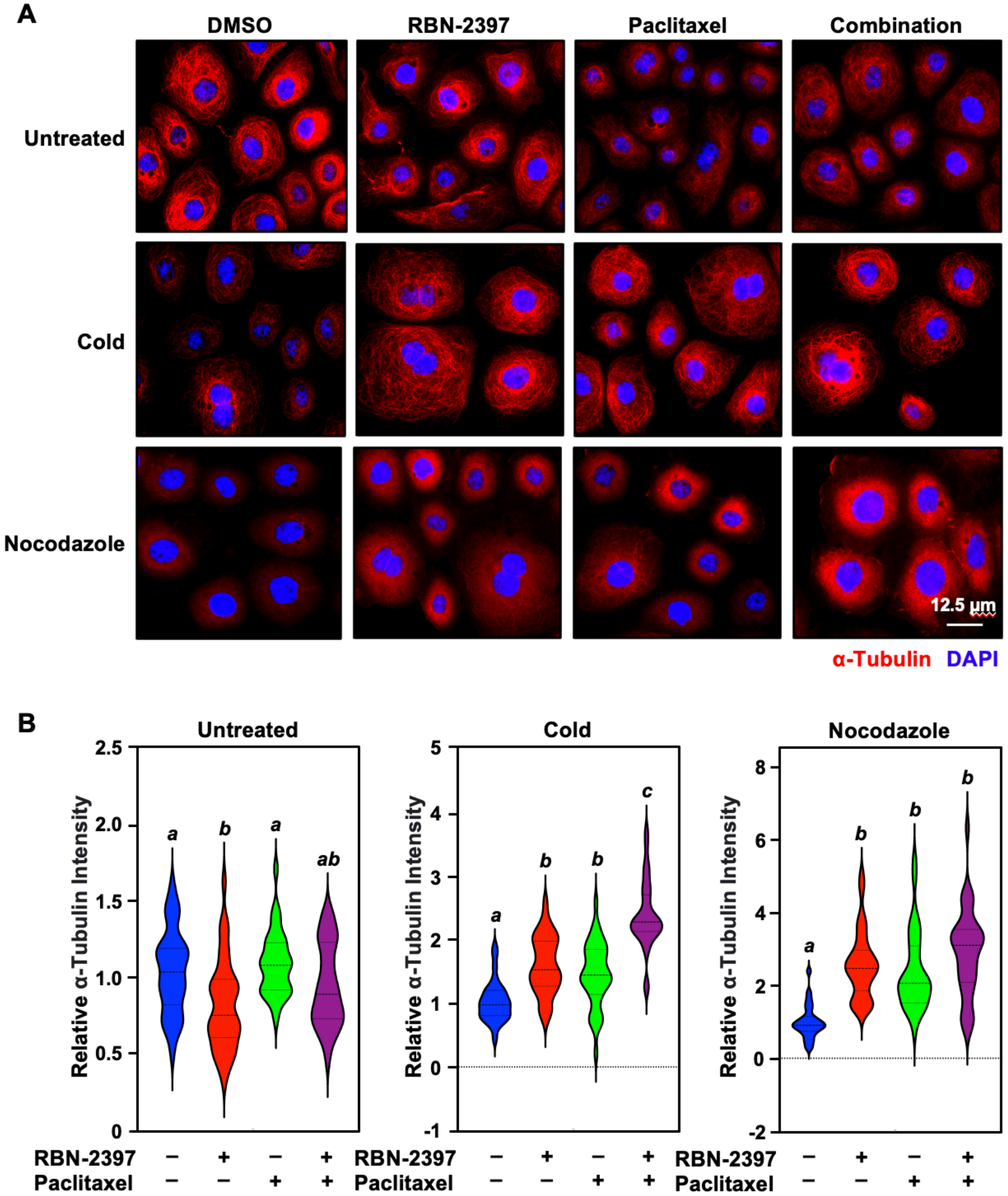
A
DMSO
RBN-2397
Paclitaxel
Combination
Untreated
Cold
Nocodazole
12.5 µm
α-Tubulin
DAPI
B
Untreated
Cold
Nocodazole
Relative α-Tubulin Intensity
a
b
a
ab
a
b
b
c
a
b
b
b
RBN-2397
Paclitaxel

**Supplementary Figure 4. Effect of RBN-2397, paclitaxel, and their combination on ovarian cancer cell biology.**

**(A and C)** Venn diagrams depicting differentially expressed genes from RNA-seq in OVCAR4 (A) and OVCAR3 (C) cells treated with RBN-2397 (*red*; 500 nM), paclitaxel (*green*; 0.4 nM) or combination (*blue*).

**(B and D)** Gene ontology analysis for genes up- or downregulated in combination treatment in OVCAR4 (B) and OVCAR3 (D) cells. The percentage of targets and p-value for each term are shown. Analysis was conducted using the Database for Annotation, Visualization, and Integrated Discovery (DAVID) Bioinformatics Resources tool.

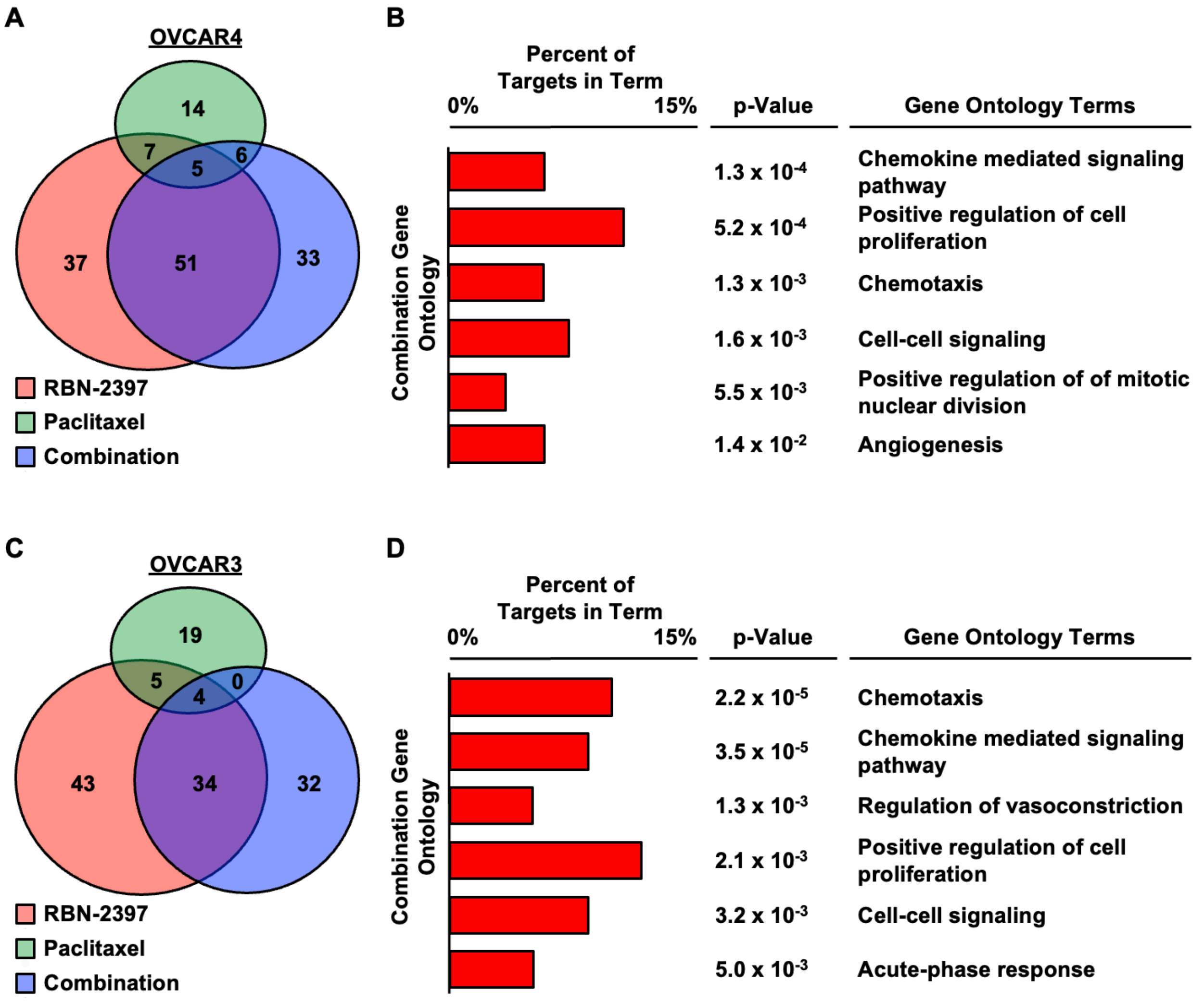
